# Dendrite-specific synaptic inputs onto dopaminergic SNc neurons

**DOI:** 10.64898/2026.08.02.742307

**Authors:** RC Evans, R Zhang, ZM Khaliq

## Abstract

Substantia nigra pars compacta (SNc) dopaminergic neurons integrate synaptic inputs through dendrites that project laterally within the SNc and ventrally into the substantia nigra pars reticulata (SNr), which then determines dopaminergic output. We use spatially-specific optogenetics to map excitatory subthalamic nucleus (STN) and pedunculopontine nucleus (PPN) as well as inhibitory SNr inputs that provide feedforward inhibition onto SNc neurons. Using multiple GABAergic mouse lines, we find that SNr inputs selectively inhibit SNc somas and proximal dendrites without preference for SNr versus SNc dendrites. Using glutamatergic mouse lines, we find that PPN inputs selectively excite somas and proximal dendrites, while STN inputs selectively excite SNr dendrites, mirroring dendrite-selective inhibition from striosomes. These findings reveal a high level of spatial organization in SNc synaptic inputs and point to separate functional roles for STN versus PPN control of dopaminergic activity.

**Graphical Abstract:** 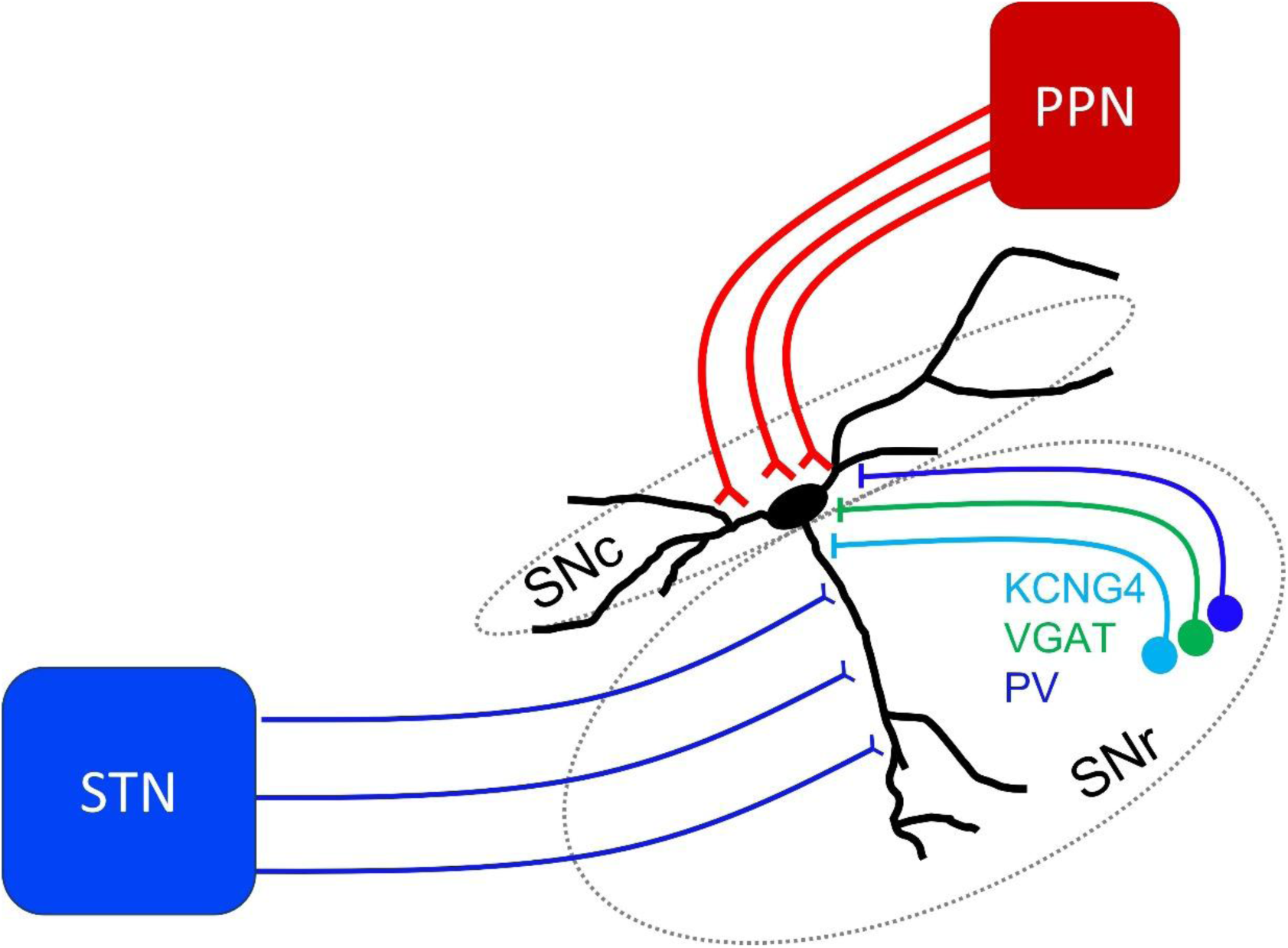

## Introduction

Midbrain dopaminergic neurons are critical for reward processing, learning, and motor control (Berke, 2018; Carmichael et al., 2021; Lerner et al., 2021; Watabe-Uchida et al., 2017). To perform these functions, dopaminergic neurons integrate excitatory and inhibitory inputs from many brain structures involved in sensory processing and motor control (Lerner et al., 2015; Watabe-Uchida et al., 2012; Wu et al., 2019). Dopaminergic neurons process diverse information, despite having a relatively simple dendritic morphology (Evans et al., 2020; Grace and Bunney, 1983), and sparse dendritic spines (Hage et al., 2016; Henny et al., 2012). In spiny neurons with more complex dendritic arbors, the location of inhibitory and excitatory synapses on somatodendritic compartments strongly controls their functional impact (Bar-Ilan et al., 2012; Boivin and Nedivi, 2018; Gidon and Segev, 2012; Liu, 2004). Understanding the fundamental organization and function of excitatory and inhibitory synaptic inputs along the somatodendritic structure of dopaminergic neurons will be critical for dissecting mechanisms of circuit level integration and the contribution of SNc inputs to reward and motor-learning.

Substantia nigra pars compacta (SNc) dopaminergic neurons have two distinct types of dendrites. Dendrites that project along the SNc, called “SNc dendrites” and dendrites that project down into the GABAergic Substantia nigra pars reticulata (SNr) nucleus called “SNr dendrites” (Björklund and Lindvall, 1975; Cheramy et al., 1981; Grace and Bunney, 1983; Henny et al., 2012; Juraska et al., 1977; Wassef et al., 1981). These dendrite types are morphologically and functionally distinct (Evans et al., 2020). The SNr dendrites are long with minimal branching and selectively receive inhibitory input from striosomal direct pathway striatal spiny projection neurons, clustering together in striosome-dendron bouquets (Crittenden et al., 2016; Evans et al., 2020). SNc dendrites, on the other hand, may be specialized for excitatory synaptic input.

The axon initial segment preferentially originates from SNc dendrites rather than directly from the somas or SNr dendrites (Evans et al., 2020). Because axon-bearing dendrites have a strong influence over action potential activity in SNc neurons, it is important to understand whether excitatory inputs synapse preferentially on the SNc vs SNr dendrites. Similarly, local dendritic interactions play an important role in the ability of inhibitory input to interrupt excitatory input (Boivin and Nedivi, 2018; Gidon and Segev, 2012); therefore, excitatory synaptic inputs selective to the SNr dendrites would preferentially interact with inhibitory input from the striatum.

Here we map three sources of input onto dopaminergic SNc neurons and evaluate their innervation of the SNc and SNr dendrites. We find that GABAergic neurons in the SNr, known to innervate SNc dopaminergic neurons (Ambrosi and Lerner, 2022; Rizzi and Tan, 2019; Tepper et al., 1995), and previously hypothesized to innervate SNr dendrites (Hajós and Greenfield, 1994), surprisingly selectively innervate dopaminergic somas and proximal dendrites. Second, we compare excitatory input from the subthalamic nucleus (STN) and the pedunculopontine nucleus (PPN), structures that excite dopaminergic neurons (Beaudoin et al., 2018; Futami et al., 1995; Galtieri et al., 2017). We find that the STN selectively excites dopaminergic SNr dendrites and is capable of evoking T-type calcium channel-mediated dendritic events, while the PPN preferentially excites dopaminergic somas and proximal dendrites. Together, these findings predict a highly selective interaction between STN excitatory input and striatal inhibitory input on dopaminergic SNr dendrites.

## Results

### Multiple SNr GABAergic subpopulations inhibit SNc dopaminergic neurons

An outstanding question is how distinct substantia nigra pars reticulata (SNr) GABAergic subpopulations differentially target SNc dopaminergic (DA) neurons, particularly regarding the spatial organization of their synaptic inputs onto SNc cells. SNr inhibition of dopaminergic SNc neurons is well-documented and is a critical component of basal ganglia circuitry (Ambrosi and Lerner, 2022; Rizzi and Tan, 2019; Tepper et al., 1995). However, the SNr GABAergic neurons are heterogeneous, with distinct genetically-, anatomically-, and behaviorally-defined SNr subpopulations identified and characterized (Deniau and Chevalier, 1992; Falasconi et al., 2025; Lee and Tepper, 2007; Liu et al., 2020; McElvain et al., 2021; Partanen and Achim, 2022; Rizzi and Tan, 2019). Parvalbumin (PV) labels a laterally-located population of SNr GABAergic neurons (Gerfen et al., 1985), that can be functionally separated from the GAD2-positive (Liu et al., 2020) and the VGAT-positive (Rizzi and Tan, 2019) populations. In addition, we used a novel SNr population marker, KCNG4, the Kv6.4 potassium channel modifying subunit. This marker has not previously been used to label SNr neurons, but co-localizes with PV-positive neurons in other structures such as the globus pallidus externa (Pamukcu et al., 2020) and cortex (Ganesh et al., 2026). Thus, we set out to systematically map how these subpopulations differentially innervate dendrites of SNc DA neurons.

To characterize the input from each GABAergic SNr subpopulation onto SNc dopaminergic neurons, we injected a Cre-dependent channelrhodopsin (CoChR) into the substantia nigra of VGAT-Cre, PV-Cre, or KCNG4-Cre mice. After 3-4 weeks to allow for viral expression, we prepared midbrain slices for electrophysiology. Recordings were made at 30-34°C and in the presence of glutamatergic receptor blockers (AP5 50 μM and either NBQX 5 μM or CNQX 12.5 μM). SNc dopaminergic neurons were recorded in whole-cell voltage clamp with cells held at - 50mV. SNr axons of each subpopulation were optically stimulated using a blue LED (473 nm) and full-field illumination. Optically-evoked inhibitory post-synaptic currents (oIPSCs) were recorded in response to a 2 second, 20 Hz train of 2 ms wide optical pulses.

We found that activating VGAT-positive SNr neurons evoked larger IPSCs than activating either KCNG4- or PV-positive SNr neurons (Figure 1B&C). Interestingly, short-term synaptic plasticity characteristics were similar across populations, depressing to 60-80% of the initial current amplitude over the course of the 2 second stimulation train (Figure 1D). Because KCNG4-positive neurons often co-express PV (Ganesh et al., 2026; Pamukcu et al., 2020), we ran an *in situ* hybridization assay (RNAscope) for PV and KCNG4. We found near perfect overlap in these populations (Figure 1E), indicating that PV and KCNG4 label the same SNr subpopulation. Comparing the amplitude of the first oIPSC in the stimulation train (Figure B, inset), we found that the evoked current amplitude was significantly larger when the VGAT SNr population was activated compared to the PV/KCNG4 SNr population (PV/KCNG4, mean oIPSC amplitude = 49.6 ± 13.9 pA, n=9, N=3; VGAT, mean oIPSC amplitude = 150.9 ± 23.5 pA, n=11, N=4; 2-tailed Mann Whitney U=13, p=0.004). These data show that multiple subpopulations of SNr GABAergic neurons inhibit SNc dopaminergic neurons, but that the VGAT-positive subpopulation exerts the strongest inhibitory effect.

**Figure 1.**
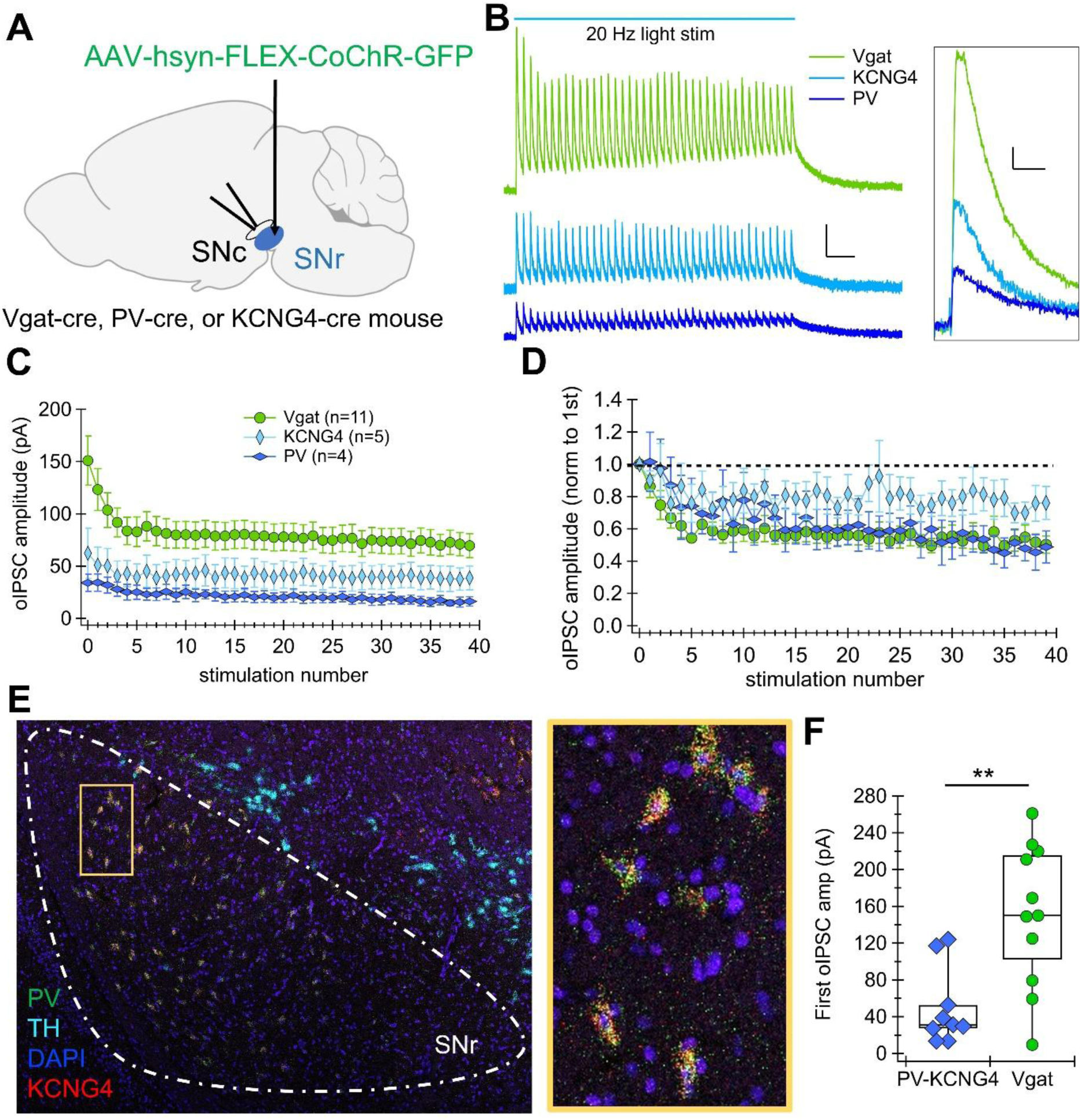
Inhibitory input from subpopulations of SNr neurons to SNc dopaminergic neurons. **A.** Schematic showing injection into SNr and recording from a SNc dopaminergic neuron. **B.** Example optically-evoked inhibitory post-synaptic currents (oIPSCs) recorded in SNc dopaminergic neurons while stimulating three genetically-defined SNr subpopulations: PV=parvalbumin-positive, KCNG4-potassium voltage-gated channel modifier Kv6.4-positive, VGAT=vesicular GABA transporter-positive. Scale bars: 50 pA, 200 ms (left); 20 pA, 10 ms (right). **C.** Averaged oIPSC current amplitude over 40 stimulations (20 Hz). **D.** Same current amplitudes as in *C*, but normalized to the amplitude of the first current. **E.** *In situ* hybridization showing Parvalbumin (PV, green), KCNG4 (red), tyrosine hydroxylase (TH, cyan), and nuclei (DAPI, blue) in a midbrain slice. Inset shows close up of location outlined in yellow. Note KCNG4 and PV show extensive overlap. **F.** Amplitude of the first oIPSC in the train is larger when activating Vgat SNr neurons vs. activating PV-KCNG4 SNr neurons. **p<0.005

### SNr inhibitory input onto SNc dopaminergic neurons occurs at the soma and proximal dendrites

SNc dopaminergic neurons have two distinct dendrite types: dendrites that follow the SNc cell body layer (“SNc dendrites”) and dendrites that project ventrally into the SNr (“SNr dendrites”) (Björklund and Lindvall, 1975; Cheramy et al., 1981; Henny et al., 2012; Wassef et al., 1981). The SNr dendrites receive specialized inhibitory input from the striatum (Crittenden et al., 2016; Evans, 2022; Evans et al., 2020), and it has been hypothesized that these dendrites would also receive strong inhibitory input from the SNr GABAergic neurons themselves (Hajós and Greenfield, 1994). However, morphological reconstructions show that SNr GABAergic axons branch extensively within the SNc cell body layer, suggesting SNc somas may be the primary target of SNr inhibition (Cebrián et al., 2005; Mailly et al., 2003).

To test these competing hypotheses and determine where along the somatodendritic domain the SNr functionally inhibits SNc neurons, we functionally mapped the strength of inhibitory synaptic inputs along SNr and SNc dendrites. After injecting a Cre-dependent CoChR into the SNr of VGAT-Cre, PV-Cre, or KCNG4-Cre mice, we used spatially-specific (∼20-30 µm diameter of CoChR activation) optogenetic stimulation (Evans et al., 2020) to selectively activate SNr axons at distinct dendritic locations (Figure 2A). To prevent action potential propagation throughout the stimulated SNr axons, we added TTX (0.5 μM) and 4-AP (300 μM) to the recording solution to block sodium and potassium channels, respectively. We measured oIPSCs in response to localized laser stimulation (5 pulses, 20 Hz) at each location. We found that oIPSCs were largest when SNr axons were activated at the soma and proximal dendrites of SNc DA neurons (Figure 2). This location selectivity was consistent across all SNr populations stimulated (PV, KCNG4, & VGAT, Figure 2D). Overall, these data show that the SNr inhibits SNc dopaminergic neurons most strongly on somas and proximal SNr and SNc dendrites.

**Figure 2.**
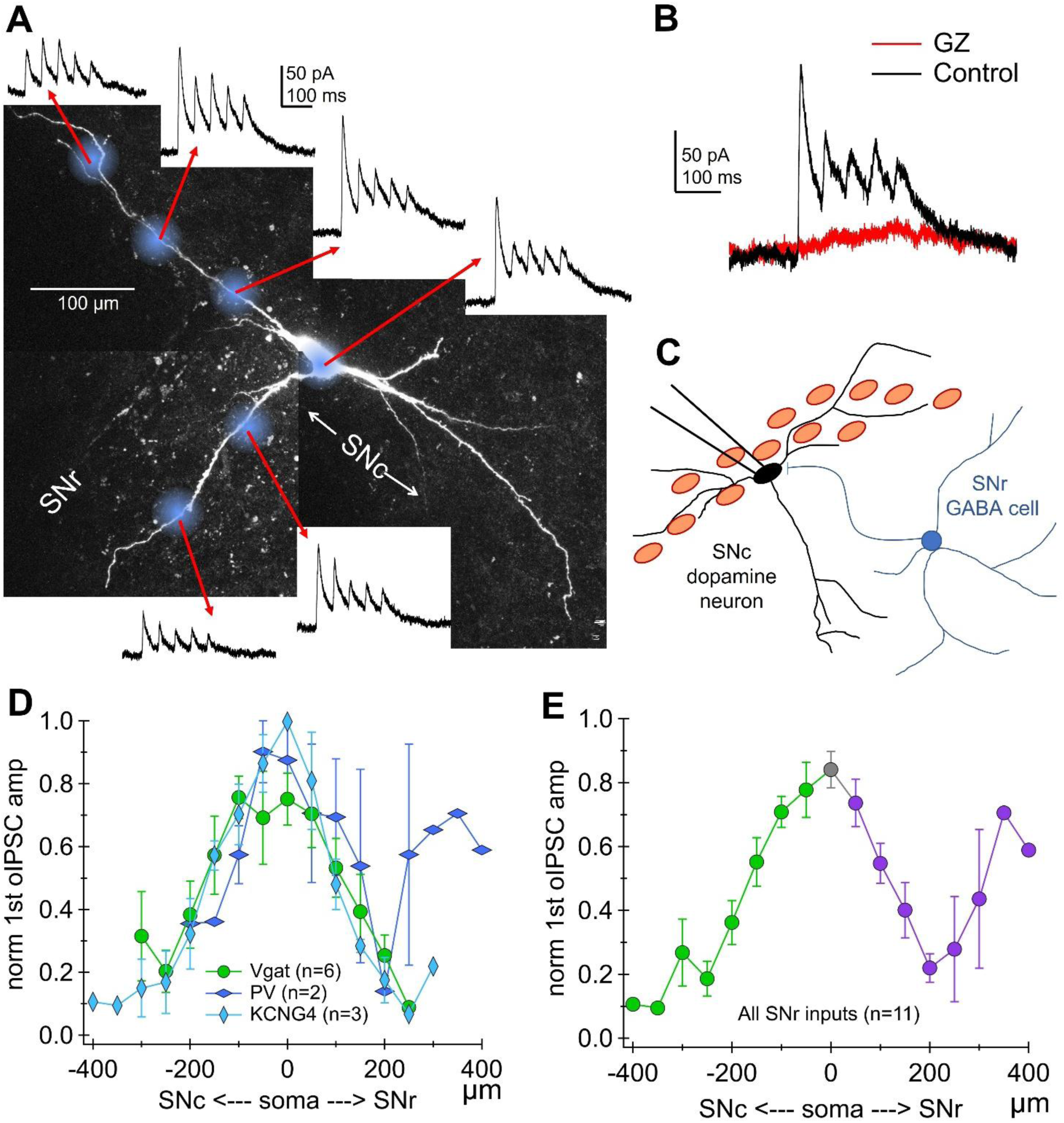
The SNr primarily inhibits SNc dopaminergic neurons on the soma and proximal dendrites. **A.**2-photon image of filled SNc neuron. Blue spots indicate locations of focal optogenetic activation of SNr axons. Example inhibitory currents recorded while stimulating axons at each location. **B.**Synaptic currents in control conditions and with GABA-A receptors blocked by gabazine (GZ). **C.** Schematic of SNr input onto SNc soma. **D.** Normalized amplitude of first optically-induces inhibitory post-synaptic current (oIPSC) (fraction of maximal) plotted against distance from the soma along the SNr (right) and SNc (left) dendrite. **E.**Same data as D, but consolidated across SNr subpopulation types.

### PPN excitation of the SNc is stronger than STN excitation

Both the pedunculopontine nucleus (PPN) and the subthalamic nucleus (STN) make monosynaptic excitatory connections onto SNc dopaminergic neurons (Watabe-Uchida et al., 2012). To compare the synaptic characteristics between PPN and STN inputs to SNc neurons, we injected a Cre-dependent CoChR-GFP into the PPN of VGluT2-Cre mice or the STN of KCNG4-Cre mice (Figure 3A&C). While VGluT2-Cre mouse line shows wide expression in regions surrounding the STN such as the lateral hypothalamus, KCNG4-Cre positive neurons are restricted mainly to STN making this mouse line preferable for our experiments.

Imaging the GFP-labeled axons of the PPN and STN inputs to the SNc revealed strikingly different axon distribution patterns. PPN axons filled the SNc cell body layer and did not strongly innervate the SNr (Figure 3B). By contrast, KCNG4-positive STN axons avoided the SNc cell body layer, preferentially targeting the SNr (Figure 3D). To measure the synaptic currents evoked by each input, we recorded SNc dopaminergic neurons in whole-cell voltage clamp, holding at -70 mV while optically-stimulating either PPN or STN axons using a full-field blue light stimulus (473 nm LED). Recordings were made in the presence of gabazine (GZ, 10 μM) and CGP55845 (1 μM) to block GABA-A and GABA-B receptors respectively. Optically-evoked excitatory post-synaptic currents (oEPSCs) were recorded in response to a 2 second, 20 Hz train of 2 ms wide optical pulses. We found that the PPN evoked large oEPSCs that depressed to 40% of their initial amplitude over the course of the stimulation train, while STN stimulation evoked small synaptic currents that slightly facilitated over the course of the stimulation train (Figures 3E-G). Comparing the first oEPSC in the train, we found that synaptic currents evoked by PPN stimulation was significantly larger than those evoked by STN stimulation (PPN -313.9 ± 84 pA n=9; STN -11.5 ± 3.7 pA n= 6; 2-tailed t-test, t=-3.59, p=0.007). These findings are consistent with previous work testing PPN and STN connectivity with SNc dopaminergic neurons (Futami et al., 1995; Galtieri et al., 2017; Iribe et al., 1999), and showing larger synaptic currents when stimulating CaMKIIa-positive PPN vs STN axons (Beaudoin et al., 2018). Therefore, these experiments show that KCNG4 labels an SNc-projecting population of STN neurons, and that the PPN input to the SNc is stronger than this STN input.

### PPN glutamatergic neurons preferentially excite SNc somas and proximal dendrites

To determine the dendritic location of PPN excitatory synaptic input, we mapped excitatory synaptic strength along both the SNr and SNc dendrites of SNc dopaminergic neurons. After injecting a Cre-dependent CoChR into the PPN of Vglut2-Cre mice, we used spatially-specific optogenetic stimulation to selectively activate PPN axons at distinct dendritic locations (Figure 3H). As described above, we added TTX (0.5 μM) and 4-AP (300 μM), and measured oEPSCs in response to localized laser stimulation (5 pulses, 20 Hz) at each location (the first current is shown in Figure 3). PPN-activated oEPSCs were reliably measured from stimulation of inputs onto the soma and proximal dendrites. However, the current amplitude sharply decayed with increased distance from the soma along both the SNr and SNc dendrites (Figure 3I). Interestingly, a slight bias toward larger currents on the SNc dendrite appears at 100 μm from the soma. However, this difference was not statistically significant (oEPSC amplitude SNc dend 80 ± 22 pA, n=8; SNr dend 53 ± 25 pA, n=6; 2-tailed Mann Whitney, U=32, p=0.172). Together these data show that PPN glutamatergic input to the SNc is strongest at the soma and proximal dendrite with no significant preference for SNc vs SNr dendrite.

**Figure 3.**
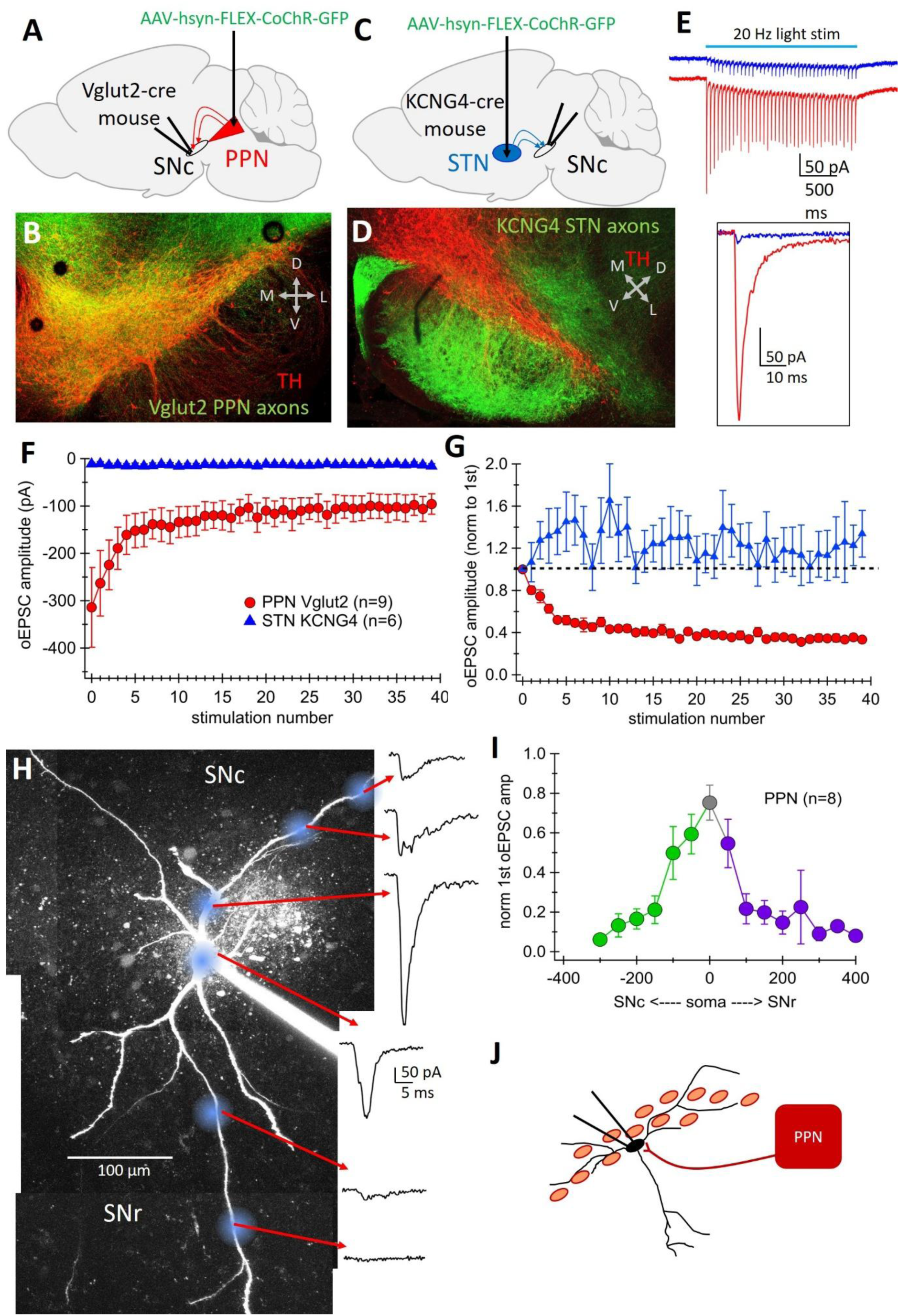
PPN excitatory input onto SNc dopaminergic neurons is stronger than STN input. **A.** Schematic of experiment. Cre-dependent CoChR injected into the PPN of Vglut2-Cre mice. Electrophysiology performed in the SNc. **B.** Confocal image of PPN glutamatergic axons (*green*) and tyrosine hydroxylase (TH, *red*) immunostaining. Note the extensive overlap between PPN axons and SNc cell body layer. **C.** Schematic for STN stimulation experiments. Cre-dependent CoChR injected into STN of KCNG4-Cre mice. Electrophysiology performed in the SNc. **D.** KCNG4-positive STN axons (*green*) and TH (*red*), not prominent STN axon expression in the SNr, ventral to the SNc. **E.** Example voltage clamp recordings from SNc neurons when optically activating PPN (*red*) or STN (*blue*) axons with a 20 Hz train of blue light. Inset: close-up of first optically-evoked excitatory post-synaptic current (oEPSC) in the train. **F.** Amplitude of each oEPSC during the 20Hz train averaged over cells. Note larger currents when PPN axons are activated. **G.** Normalized oEPSC amplitude (fraction of first current amplitude) during the 20 Hz train. **H.** Filled SNc dopaminergic neuron. Blue spots indicate locations of focal optogenetic activation of PPN axons. Example excitatory currents recorded while stimulating axons at each location. **I.** Amplitude of first oEPSC normalized to maximum current plotted against distance from the soma along the SNr (*right*) and SNc (*left*) dendrite. **J.** Schematic of PPN input onto SNc soma.

### The STN preferentially excites the SNr dendrite of dopaminergic neurons

We have previously found that an inhibitory input from the striatal striosome compartments selectively inhibits the SNr dendrites of SNc dopaminergic neurons (Evans et al., 2020). However, a parallel excitatory input selectively onto these dendrites has not been identified. To determine whether the STN selectively excites one SNc somatodendritic domain, we used the same spatial optogenetic approach described above while activating KCNG4-positive STN axons. We found that STN-evoked currents were reliably recorded at the soma of SNc dopaminergic neurons, and that the current amplitude dropped sharply along the SNc, but not the SNr dendrite. In some cells, stimulating STN axons on the SNr dendrite evoked even larger oEPSCs than at the soma (Figure 4A). These findings show a functional correlate to the anatomical innervation pattern observed with confocal imaging, where STN axons avoid the SNc cell body layer but thoroughly fill the SNr (Figure 3D). Indeed, we find that the relative current amplitude generated by STN axon activation is maintained along the whole SNr dendrite up to at least 300 μm distant from the soma (Figure 4C). By contrast, synaptic current amplitude is greatly attenuated even 50 μm away from the soma along the SNc dendrite. These data show remarkable dendrite selectivity of STN excitatory synaptic inputs for the ventrally-projecting SNr dendrites of SNc dopaminergic neurons.

Extensive previous work has shown that dendrite-specific excitation can cause large isolated dendritic events, called dendritic spikes, in pyramidal and (Stuart and Spruston, 2015) striatal spiny (Plotkin et al., 2011) neurons. However, it is not clear that dendrite-selective excitatory input onto dopaminergic neurons could induce such events. Interestingly, the example neuron shown in Figure 4D exhibits a large slow inward current when STN axons were activated at the most distal SNr dendritic regions (Figure 4D). To determine whether this slow inward current was due to dendritic calcium channels, we washed on the T-type calcium channel blocker, TTA-P2 (1 μM). Blocking T-type calcium channels eliminated the large slow inward current, leaving synaptic inward currents intact (Figure 4D). We observed this behavior rarely (1 out of 6 neurons), however this suggests that STN inuputs can interact with dendritic calcium channels, such as T-type channels, to modulate dendritic excitability and tune cellular responses to synaptic input.

**Figure 4.**
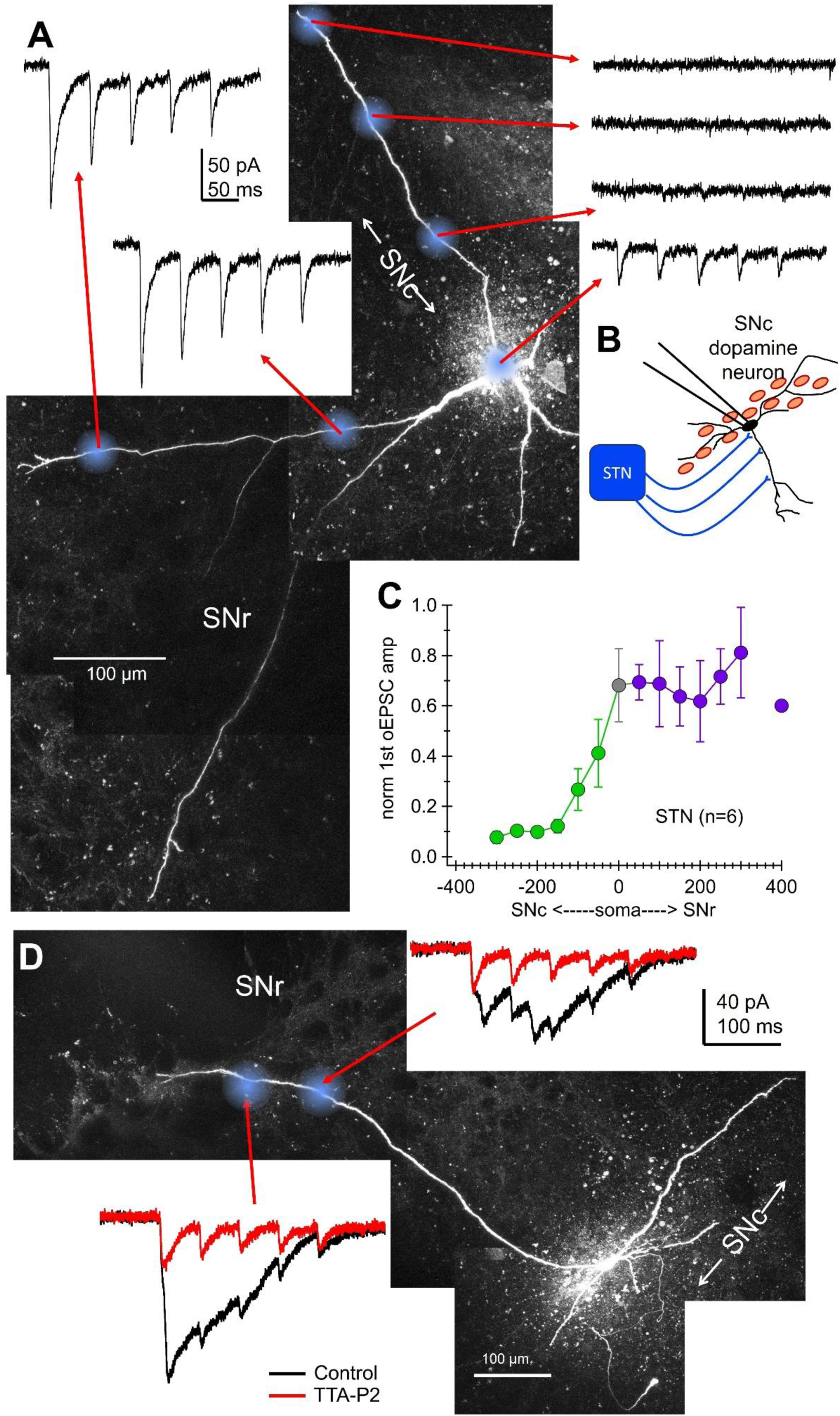
STN inputs primarily excite the SNr dendrite. **A.** Filled SNc dopaminergic neuron. Blue spots indicate locations of focal optogenetic activation of KCNG4-positive STN axons. Example excitatory currents recorded while stimulating axons at each location. **B.** Schematic of STN functional excitatory synapse location. **C.** Normalized amplitude of first optically-induced inhibitory post-synaptic current (oIPSC) (fraction of maximal) plotted against distance from the soma along the SNr (right) and SNc (left) dendrite. **D.** STN input on the most distal SNr dendrites can evoke T-type calcium channel mediated dendritic events. Filled SNc neuron. Spatially-specific STN axon activation on the distal SNr dendrite evoked dendritic-spike-like events that were eliminated by T-type calcium channel blockers (TTA-P2).

## Discussion

Here we mapped excitatory and inhibitory inputs along distinct dendrite types (SNr and SNc dendrites) of SNc dopaminergic neurons. While early studies suggested that SNr GABAergic neurons primarily innervate ventrally-projecting “SNr dendrites”, our optical mapping results demonstrate that SNr inhibition is primarily targeted to the perisomatic region of the SNc neurons. Similarly, we find that the PPN preferentially excites SNc somas and proximal dendrites with no significant bias for dendrite type. By contrast, STN inputs are highly selective for SNr dendrites over SNc dendrites. When combined with previous dendritic mapping (Evans et al., 2020), it is clear that excitatory input from the STN mirrors inhibition from striatal striosomes, which also selectively innervate dopaminergic SNr dendrites. On the other hand, excitatory input from the PPN is locally opposed by inhibition from the SNr and globus pallidus externa (GPe), which both selectively innervate SNc somas and proximal dendrites. Thus, these findings demonstrate functionally distinct dendrite types in SNc neurons, and reveal organizational principles co-localizing excitation and inhibition from specific nuclei.

### STN selective excitation of the SNr dendrite

The location of inhibitory and excitatory synapses on somatodendritic compartments can govern their functional impact (Bar-Ilan et al., 2012; Boivin and Nedivi, 2018; Gidon and Segev, 2012; Liu, 2004). Dendrite-selective excitation often functions to generate dendritic calcium spikes (Stuart and Spruston, 2015) or neuronal up-states (Plotkin et al., 2011), and to promote synaptic plasticity (Remy and Spruston, 2007). However, dendrite-selective inputs have been predominately studied in hippocampal and cortical pyramidal neurons, which have a high density of dendritic spines, low membrane resistances, and strikingly divergent apical and basal dendritic structures. The SNc dopaminergic neurons, on the other hand, have sparse spines (Hage et al., 2016), are electrotonically compact (Gentet and Williams, 2007; Hage and Khaliq, 2015; Häusser et al., 1995), and have SNr and SNc dendrites with minimal branching and relatively small differences in dendritic structure (Evans et al., 2020; Henny et al., 2012; Montero et al., 2021). Our previous work shows that the SNr dendrite is under powerful inhibitory control from striosome compartments of the striatum (Evans et al., 2020) whose axons fully surround SNr-projecting dendrites forming striosome-dendron bouquets (Crittenden et al., 2016). The co-localization of STN excitation and striosomal inhibition on the SNr dendrite suggests that these two inputs to the SNc directly oppose each other (Supplemental Figure S1). Interestingly, the striosomal inhibition of the SNr dendrite is surprisingly strong for a dendrite-specific inhibition, while the STN generates very small excitatory currents. This suggests that rather than driving action potential firing, the primary role of STN input may be to modulate the responsivity of SNc neurons to other inputs. Indeed, slight synaptically-evoked depolarization can dramatically increase NMDA receptor-mediated calcium influx in SNc dendrites (Hage et al., 2016; Hage and Khaliq, 2015). Therefore, STN inputs may serve to modulate the gain of inputs from other neurons in an NMDA-dependent manner. Similarly, because STN input to the SNc has a significant NMDA-mediated component (Beaudoin et al., 2018), inputs from other neurons may augment the impact of STN excitation.

Two additional possible functions for the STN’s weak, dendrite-specific excitatory input to the SNc could be 1) to directly promote dendritic dopamine release into the SNr (Beckstead et al., 2004; Bulumulla et al., 2022; Gantz et al., 2013; Hikima et al., 2021; Zhou et al., 2009), or 2) to act synergistically with the release of inhibition to amplify SNc T-type mediated rebound activity (Beaver and Evans, 2025; Evans, 2022; Evans et al., 2017). Because striosomal input greatly facilitates T-type mediated rebound activity, the release of striosomal inhibition at the same time as STN excitation, could be optimized for evoking rebound burst activity in the SNc neuron. Future work is needed to understand network interactions that would control the physiological timing between striosomal inhibition and STN excitation of the same dendrite.

Finally, it is possible that STN modulation of the SNc is predominately mediated by feed-forward inhibition through the SNr. Recent work shows that *in vivo* stimulation of STN inputs actually inhibit a population of SNc neurons (Hadjas et al., 2025), and *ex vivo* recordings have shown STN-evoked inhibitory currents can occur on both SNr (Sitzia et al., 2024) and SNc (Iribe et al., 1999) neurons. Interestingly, STN feed-forward inhibition could serve to electrically isolate the SNr dendrite because STN activation of SNr GABAergic neurons would inhibit SNc somas while the STN selectively excites the SNr dendrite. Future work measuring dendritic calcium during STN stimulation of the SNc is needed to determine the extent to which isolated dendritic calcium events could be generated under these conditions.

### Synaptic interactions on the soma and proximal dendrites

We found that PPN input to SNc neurons was proximal to the soma and decays similarly along the SNr and SNc dendrite. Previous work has shown that a low percentage of SNc neurons respond to PPN input onto dendrites more distal than ∼80 µm from the soma, while a high percentage of SNc neurons respond to STN stimulation along dendrites as far as 120 µm from the soma (Galtieri et al., 2017). However, this study did not distinguish between SNr vs SNc dendrites. Here we show that STN axon activation generates large synaptic currents as far as 300 µm from the soma, but only along the SNr dendrite. PPN inputs have been shown to strongly drive action potentials in SNc neurons (Galtieri et al., 2017). This could relate to the relative location of PPN synapses near the axon initial segment. SNc axons originate from dendrites rather than somas (Grace and Bunney, 1983; Häusser et al., 1995) and specifically originate from SNc dendrites (Evans et al., 2020). Because excitatory synapses on axon-bearing dendrites are in a privileged position with regard to action potential generation (Thome et al., 2014), and synaptic excitation onto proximal dendrites effectively generates action potentials (Blythe et al., 2009), PPN input is positioned to directly control the action potential output of SNc neurons.

Past studies showed that dendrites of dopaminergic neurons located in the SNr receive a greater density of GABAergic synapses relative to dendrites located within the SNc (Henny et al., 2012), raising the question of whether SNr GABAergic inhibition primarily targets this dendrite. Axonal reconstructions show SNr axons within the SNc (Cebrián et al., 2005; Mailly et al., 2003). Our functional mapping reveals that SNr inhibition of the SNc is proximal to the soma, suggesting that SNr inhibition would play a strong role in action potential suppression.

Indeed, SNr inhibition of the SNc effectively suppresses action potential output in a circuit-specific manner (Ambrosi and Lerner, 2022). This strong SNr inhibition of the SNc is controlled by the striatum, which provides inhibitory input to SNr GABAergic neurons. Striatal inhibition of SNr neurons can functionally disinhibit SNc dopaminergic neurons (Ambrosi and Lerner, 2022; Freeze et al., 2013; Tepper et al., 1995), increasing dopaminergic activity. However, striatal control of dopaminergic neurons may be highly complex due to the differential location of striatal vs SNr inhibition on SNc neurons. For example, striatal input could selectively inhibit one part of an SNc neuron (the SNr dendrite) while simultaneously disinhibiting another part (the soma). Future work is needed to determine the extent to which such compartmental isolation occurs *in vivo*.

Cognitive and motor information flow through midbrain has been proposed to involve reciprocal loops (‘ascending spiral’) that includes connections between the striatum, SNr and dopaminergic neurons (Haber et al., 2000). Ambrosi and Lerner (2022) examined these loops and found that open-loop participating neurons in the SNr produced little inhibition of DA neurons compared with closed-loop neurons. They suggested that this weak effect could be explained if open loops preferentially targeted distal dendrites of SNc DA neurons. In our study, we found that inhibitory input from SNc GABAergic neurons onto distal dendrites of dopaminergic neurons was present but resulted in substantially smaller amplitude IPSCs. Thus, our findings are consistent with the hypothesis that open loops may target distal dendrites, where inhibitory inputs produce a weaker reduction of firing. As a non-exclusive alternative, the open loops may also target perisomatic regions but produce smaller-amplitude inhibitory currents that lead to weaker effects on spiking relative to closed-loop synapses. These possibilities should be distinguished in future studies.

Our current findings along with previous mapping of basal ganglia inputs to SNc neurons suggest that interactions between inhibitory and excitatory inputs at the soma and proximal dendrites of SNc dopaminergic neurons likely play a complex role in controlling action potential firing. Specifically, our previous work shows that the globus pallidus external segment (GPe) preferentially inhibits SNc somas and proximal dendrites (Evans et al., 2020). PPN excitation, SNr inhibition, and GPe inhibition are all strong and have the potential to effectively modulate SNc neural output. The relative timing of PPN vs SNr and GPe input to the dopaminergic neurons will powerfully control action potential firing in these neurons. One additional layer of complexity comes from the connectivity between these structures. The SNr and GPe both send inhibitory inputs to the PPN (Fallah et al., 2025), and the PPN excites SNr neurons (Hadjas et al., 2025). Therefore, circuit interactions throughout the midbrain will critically determine action potential output of SNc dopaminergic neurons.

### Connections between functional subpopulations within basal ganglia nuclei

Basal ganglia nuclei usually include multiple functionally distinct subpopulations, and we have previously shown that molecularly-defined cell types within the striatum and the GPe differentially innervate SNc dopaminergic neurons (Evans et al., 2020). In the SNr, multiple functional GABAergic subpopulations have been identified (Liu et al., 2020; McElvain et al., 2021; Rizzi and Tan, 2019). The PV-positive SNr neurons were previously identified as the primary inhibitory input from the SNr to the dopaminergic neurons (Rizzi and Tan, 2019), but our results show VGAT-positive SNr neurons have a strong connection with the dopaminergic neurons in agreement with others (Ambrosi and Lerner, 2022). We see that KCNG4 and PV are overlapping populations within the SNr, as has been shown in the GPe (Pamukcu et al., 2020) and cortex (Ganesh et al., 2026). We found that all SNr populations tested primarily inhibited the soma and proximal dendrites of SNc neurons. These findings indicate that SNr input to the SNc is similar across molecularly-defined GABAergic subpopulations.

STN neurons are also functionally diverse (Prasad and Wallén-Mackenzie, 2024). The STN neurons that project to SNc dopaminergic neurons have been shown to express alpha 7 nicotinic receptors (Xiao et al., 2015), but the co-localization of KCNG4 in these neurons is not known. We used KCNG4 to define STN neurons because Vglut2 does not sufficiently separate the STN from nearby glutamatergic structures. However, KCNG4 may represent a subpopulation of STN neurons that has a specific input pattern onto SNc dopaminergic neurons, selectively exciting the SNr dendrite. We cannot exclude the possibility that a separate, KCNG4-negative population of STN neurons could innervate the somas or SNc dendrites.

Importantly, the SNc itself contains functionally distinct dopaminergic subpopulations. These SNc populations differ in their function, circuit connectivity, and vulnerability to neurodegeneration (Yamada et al., 1990; Double et al., 2010; Azcorra et al., 2023; Fushiki et al., 2024; Mantas et al., 2024; Wu et al., 2019). *In vivo*, STN and PPN excitatory input have distinct effects on these SNc populations, inhibiting vulnerable and exciting resilient SNc neurons (Hadjas et al., 2025). In the present study, ventral tier SNc neurons were targeted and most neurons (>80%) displayed the electrophysiological signature, a T-type calcium channel mediated after-depolarization, of vulnerable Aldh1a1-positive, calbindin-negative SNc neurons (Beaver and Evans, 2025; Evans et al., 2017). Future work is needed to directly compare the strength and characteristics of excitatory synaptic inputs onto vulnerable and resilient SNc neurons.

## Conclusion

Here we show that excitatory and inhibitory synaptic inputs onto SNc dopaminergic neurons are highly spatially organized onto distinct dendrite types. This dendritic organization will directly impact the function of and interactions between these synaptic inputs to control dopaminergic output.

## Resource Availability

### Lead Contact

Further information and requests for resources and reagents should be directed to and will be fulfilled by the Lead Contact Dr. Rebekah Evans.

### Materials Availability

This study did not generate unique reagents

### Data and Code Availability

Raw data will be made available upon request.

### Experimental Model and Subject Details

All animal procedures were approved by the animal care and use committee (ACUC) for the National Institute of Neurological Disorders and Stroke (NINDS) at the National Institutes of Health (NIH). Mice of both sexes were virally injected at 18 days old or greater and were used for *ex vivo* electrophysiology and two-photon imaging 3-8 weeks post-injection. The following mouse strains were used: PV-Cre (129P2-Pvalb(tm1(Cre)Abr)/J, Jax #017320), KCNG4-Cre (B6.129(SJL)-Kcng4(tm1.1(Cre)Jrs)/J, Jax #029414), VGAT-Cre (B6J.129S6(FVB)-SLc32a1(tm2(Cre)Lowl)lMwarJ, Jax #028862), and Vglut2-Cre (B6J.129S6(FVB)-SLc17a6(tm2(Cre)Lowl)lMwarJ, Jax #028863)

## Methods

### Viral injections

Viral injections were conducted on a Stoelting Stereotax. Mice were anesthetized for the duration of the stereotaxic injection with a constant flow of 1-3% isoflurane and oxygen at 1L/min. Body temperature was maintained during surgery and anesthesia recovery via a warm pad. AAV-hsyn-FLEX-CoChR (Boyden, UNC vector core) was injected bilaterally (500 nL per hemisphere) into the PPN (X: ±1.1, Y: -4.1, Z: -3.7), STN (X: ±1.5, Y: -1.8, Z: -4.7), or SNr (X: ±1.4-1.5, Y:-2.5, Z: -5.0-5.1) via Hamilton syringe. After injection was complete, the needle was raised 1-2 mm and left for 10 minutes prior to removal. In 4 of samples recorded from PPN-injected conditions, an AAV containing chrimsonR was injected into the striatum for a separate experiment that is not discussed in this paper. Because the inputs from the striatum to the SNc are inhibitory and GABA receptor blockers were present in the bath, the presence of this additional red-light activated channel rhodopsin does not significantly disrupt these experiments.

### Slicing and electrophysiology

Mice were anesthetized with isoflurane and transcardially perfused with ice cold slicing solution (modified artificial cerebrospinal fluid, ACSF) containing (in mM) 198 glycerol, 2.5 KCl, 1.2 NaH_2_PO_4_, 20 HEPES, 25 NaHCO_3_,10 glucose, 10 MgCl_2_, 0.5 CaCl_2_, 5 Na-ascorbate, 3 Na-pyruvate, and 2 thiourea. Mice were decapitated and brains extracted. Coronal slices were cut at 200 µm thickness on a vibratome and incubated for 30 minutes in heated (34°C) holding solution containing (in mM) 92 NaCl, 30 NaHCO_3_, 1.2 NaH_2_PO_4_, 2.5 KCl, 35 glucose, 20 HEPES, 2 MgCl_2_, 2 CaCl_2_, 5 Na-ascorbate, 3 Na-pyruvate, and 2 thiourea. Slices were then maintained at room temperature and used 30 min to 6 hours later. Whole-cell recordings were made using borosilicate pipettes (2-7 MΩ) filled with internal solution containing (in mM) 122 KMeSO_3_, 9 NaCl, 1.8 MgCl_2_, 4 Mg-ATP, 0.3 Na-GTP, 14 phosphocreatine, 9 HEPES, 0.45 EGTA, 0.09 CaCl_2_, 0.05 Alexa Fluor 594 hydrazide, and 0.1-0.3% neurobiotin adjusted to a pH value of 7.35 with KOH. For experiments measuring inhibitory synaptic currents, cells were voltage clamped at -50 mV. For excitatory current measurements, cells were voltage clamped at -70 mV. Cell capacitance and access resistance (< 25 MΩ) were compensated to 30-70%. Liquid junction potential (-8 mV) was not corrected. All experiments were conducted heated (31-34°C).

### Full field and spatial optogenetic activation

Experiments were conducted on an Olympus BX61W1 multiphoton upright microscope. Full field optogenetic activation of CoChR-expressing axons in brain slice was achieved by blue (470nm) LED (Thorlabs, LED4D067) light sent to the tissue via a silver mirror. Spatially-specific optogenetic experiments used a blue (473 nm) laser (Obis, Coherent) ranging from 0.6-2.7 mW measured at the back of the objective (as in Evans et al., 2020). Inhibitory currents were measured in the presence of AP5 (50 µM) and either NBQX (5 µM) or CNQX (12.5 µM), while excitatory currents were measured in the presence of gabazine (SR 95531, 10 µM) and CGP 55845 (1 µM). Spatially-specific optogenetic dendritic mapping experiments included TTX (0.5 µM) and 4-AP (300 µM).

### Fluorescence in situ hybridization (FISH)

Mouse brains were extracted and fresh frozen on dry ice in preparation for *in situ* hybridization. 16 µM thick slices were cut on a cryostat. FISH was performed using RNAscope multiplex procedure as in Evans et al., 2020. All FISH reagents are commercially available from ACD bio. Mouse parvalbumin channel 1, KCNG4 channel 2, and tyrosine hydroxylase (TH) channel 3 probes were used.

### Drugs

Salts were obtained from Millipore-Sigma. Alexa 594 (Life Technologies), TTX, gabazine, D-APV, CNQX, NBQX, and CGP 55845 were obtained from Tocris or HelloBio. TTA-P2 was obtained from Alomone. Drugs were dissolved in water except CGP 55845 and TTA-P2, which were dissolved in DMSO, and kept as concentrated stock solutions before being added to ACSF for bath application to brain slices.

### Quantification and statistical analysis

Analysis was conducted in Igor (Wavemetrics). Data normality was evaluated with the Shapiro Wilk test. If data were normally distributed, t-tests were used to compare two samples, while non-normal data were evaluated with the Mann Whitney U test. Data in text is reported as mean ± SEM and error bars in figures are ± SEM. Boxplots show medians, 25^th^ and 75^th^ percentiles (boxes) and 9^th^ and 91^st^ percentiles (whiskers). Biological replicates are individual cells and include samples from at least 3 separate mice with the exception of PV-Cre and KCNG4-Cre SNr stimulation experiments, which were recorded from 1 and 2 mice, respectively.

## Acknowledgments

Funding for this research was provided by an NINDS intramural research program grant NS003135 (to Z.M.K.) and NINDS/BRAIN Initiative K99/R00 career development grant R00NS112417 (to R.C.E.). We thank members of the Evans lab for feedback on previous versions of this manuscript.

