## Supplemental Figures for "Dendrite-specific synaptic inputs onto dopaminergic SNc neurons"

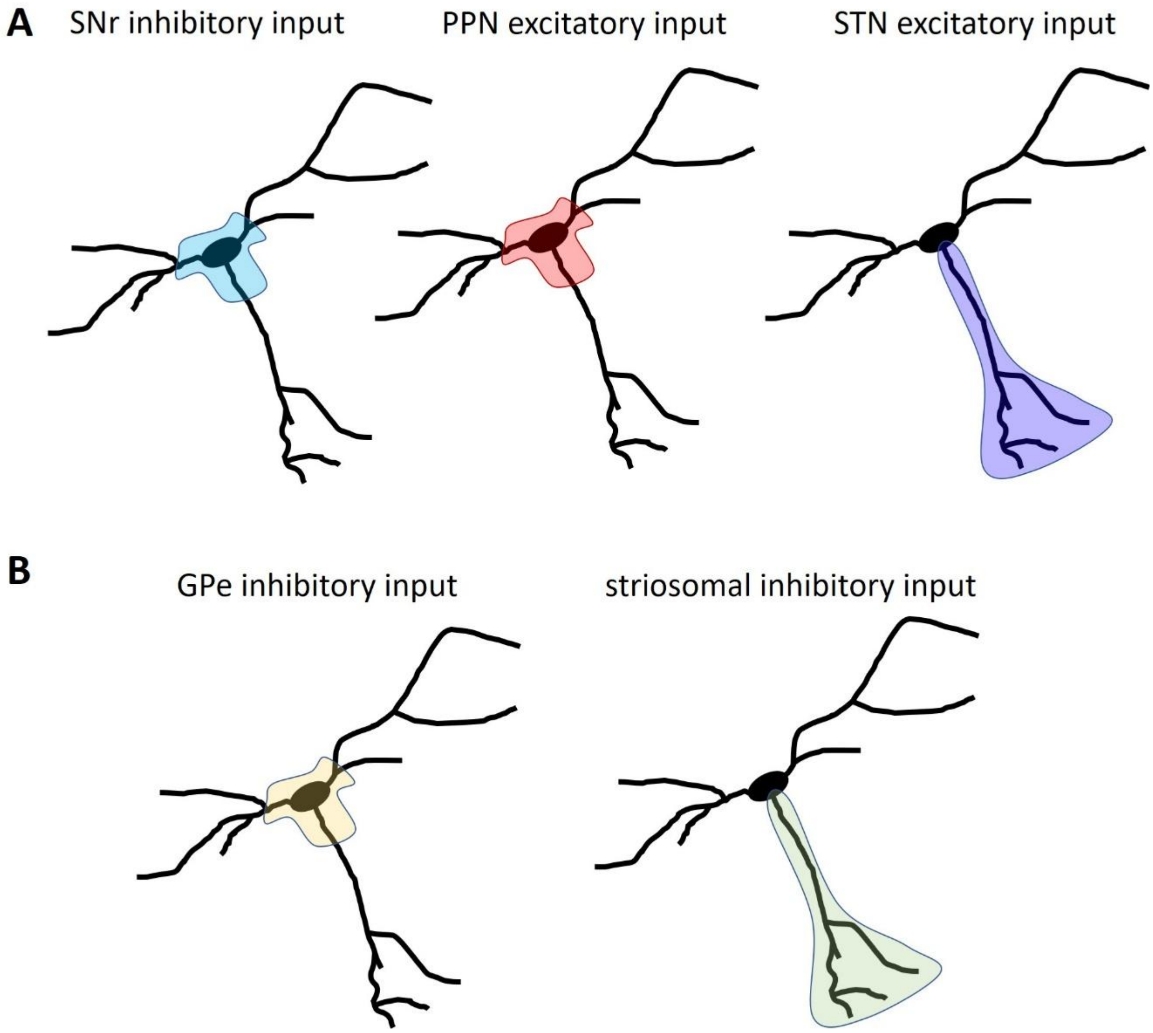


***Figure S1. Synaptic input onto SNc dopaminergic neurons is organized across distinct somatodendritic domains.*** **A.** Schematic of findings presented in this paper. The SNr and PPN preferentially synapse on the soma and proximal dendrites, while the STN selectively activates the SNc neuron's SNr dendrite. **B.** Schematic of findings from Evans et al., 2020. Globus pallidus externa (GPe) inhibitory inputs preferentially synapse on SNc somas and proximal dendrites, while striatal striosomes preferentially inhibit the SNr dendrite.
